# High-resolution screening for marine prokaryotic and eukaryotic taxa with selective preference for PE and PET surfaces

**DOI:** 10.1101/2021.12.26.474179

**Authors:** Katherine S. Marsay, Yuri Koucherov, Keren Davidov, Evgenia Iankelevich-Kounio, Sheli Itzahri, Mali Salmon-Divon, Matan Oren

## Abstract

Marine plastic debris serve as substrates for the colonization of a variety of prokaryote and eukaryote organisms. Of particular interest are the microorganisms that have adapted to thrive on plastic as they may contain genes, enzymes or pathways involved in the colonization or metabolism of plastics. We implemented DNA metabarcoding with nanopore MinION sequencing to compare the one-month-old biomes of hydrolysable (polyethylene terephthalate) and non-hydrolysable (polyethylene) plastics surfaces vs. those of glass and the surrounding water in a Mediterranean Sea marina. We sequenced longer 16S rRNA, 18S rRNA and ITS barcode loci for a more comprehensive taxonomic profiling of the bacterial, protist and fungal communities respectively. Long read sequencing enabled high-resolution mapping to genera and species. Using differential abundance screening we identified 32 bacteria and five eukaryotic species, that were differentially abundant on plastic compared to glass. This approach may be used in the future to characterize the plastisphere communities and to screen for microorganisms with a plastic-metabolism potential.

## Introduction

Marine plastic debris is a growing global pollution concern, as it jeopardizes aquatic life through entanglement, ingestion, or introduction of toxic chemicals (reviewed in (Amaral-Zettler et al., 2020)). Most plastic polymers persist for a long time in the oceans. As such, they serve as substrates for the colonization of a variety of marine organisms and the establishment of complex microorganism communities (Jacquin et al., 2019). This new human-made ecosystem is referred to as the plastisphere and includes a distinct biota from that of its surrounding waters (e.g. (Zettler et al., 2013; Bryant et al., 2016; De Tender et al., 2017)). Some marine bacteria inhabiting marine plastic debris have capabilities to degrade plastic polymers (reviewed in (Roager and Sonnenschein, 2019)) and few were shown to utilise plastics as their carbon food source (Sudhakar et al., 2008; Harshvardhan and Jha, 2013; Auta et al., 2017).

A suitable approach to investigate the assembly of the plastisphere communities is incubation experiments, where surfaces made of known materials are suspended in water for predefined time periods, either at sea (e.g. (Eich et al., 2015; Oberbeckmann et al., 2016; Pollet et al., 2018; Pinto et al., 2019; Dudek et al., 2020)) or in marine microcosms (e.g. (Ogonowski et al., 2018; Kirstein et al., 2019)). To identify taxa that preferentially colonize plastic debris, as opposed to general surface colonizers, reference substrates such as glass are often used (Oberbeckmann et al., 2016; Kirstein et al., 2018, 2019; Ogonowski et al., 2018; Pinto et al., 2019). The identification of taxa is usually done with DNA metabarcoding based on relevant genetic loci. Most plastisphere microbiome studies have used the hypervariable V3– V5 region of the 16S rRNA gene for metabarcoding of bacteria and the V4 or V9 regions of the 18S rRNA gene to identify eukaryotic taxa (reviewed in (Jacquin et al., 2019; Latva et al., 2021)). However, in most cases, the mapping of these relatively short (less than 0.5 Kbp) sequences to the databases could not resolve lower taxonomic levels and could not identify species. Additionally, the 18S barcodes and primers are limited in their coverage and do not cover certain phylogenetic groups such as fungi (Leray and Knowlton, 2016; Davidov et al., 2020). To date, only a handful of studies have assessed the fungal component of the plastisphere (De Tender et al., 2017; Kettner et al., 2017; Amend et al., 2019; Davidov et al., 2020; Lacerda et al., 2020; Yang et al., 2020; Xue et al., 2021).

We have previously established the Nanopore MinION as a tool to obtain long read sequencing that enables the identification of microbes at the species level (Davidov et al., 2020). We also demonstrated the advantages in using several complementary genetic barcodes for a more comprehensive taxonomic profiling of the plastisphere. In this study we used a similar metabarcoding approach that was based on the full 16S rRNA gene and the longer 18S V4-V5 locus, as well as the ITS2 locus, to compare between the microbial communities on polyethylene (PE), polyethylene terephthalate (PET) and glass after one-month incubation in a Mediterranean Sea marina. In continuation to our previous study, we used a modified experimental setup that included different surfaces, in five replicates, to identify taxa that were differentially abundant on plastic.

## Methods

### Experimental setup

Two 17×26 mm flat pieces of each of the three materials were used, polyethylene (PE) plastic food bags, polyethylene terephthalate (PET) drinking water bottles and glass microscope slides. Each material was tied with a fishing line to a straw and secured with plastic beads to create a ‘mobile’. Five mobiles were then secured along the dock in Herzliya Marina, Israel (32° 09′ 38.8” N 34° 47′ 35.0” E) such that the surfaces were submerged ∼0.5 m below the water surface (Fig. 1). After one month each material from the mobile was sampled for DNA extraction and metabarcoding or microscopy.

**Figure 1.**
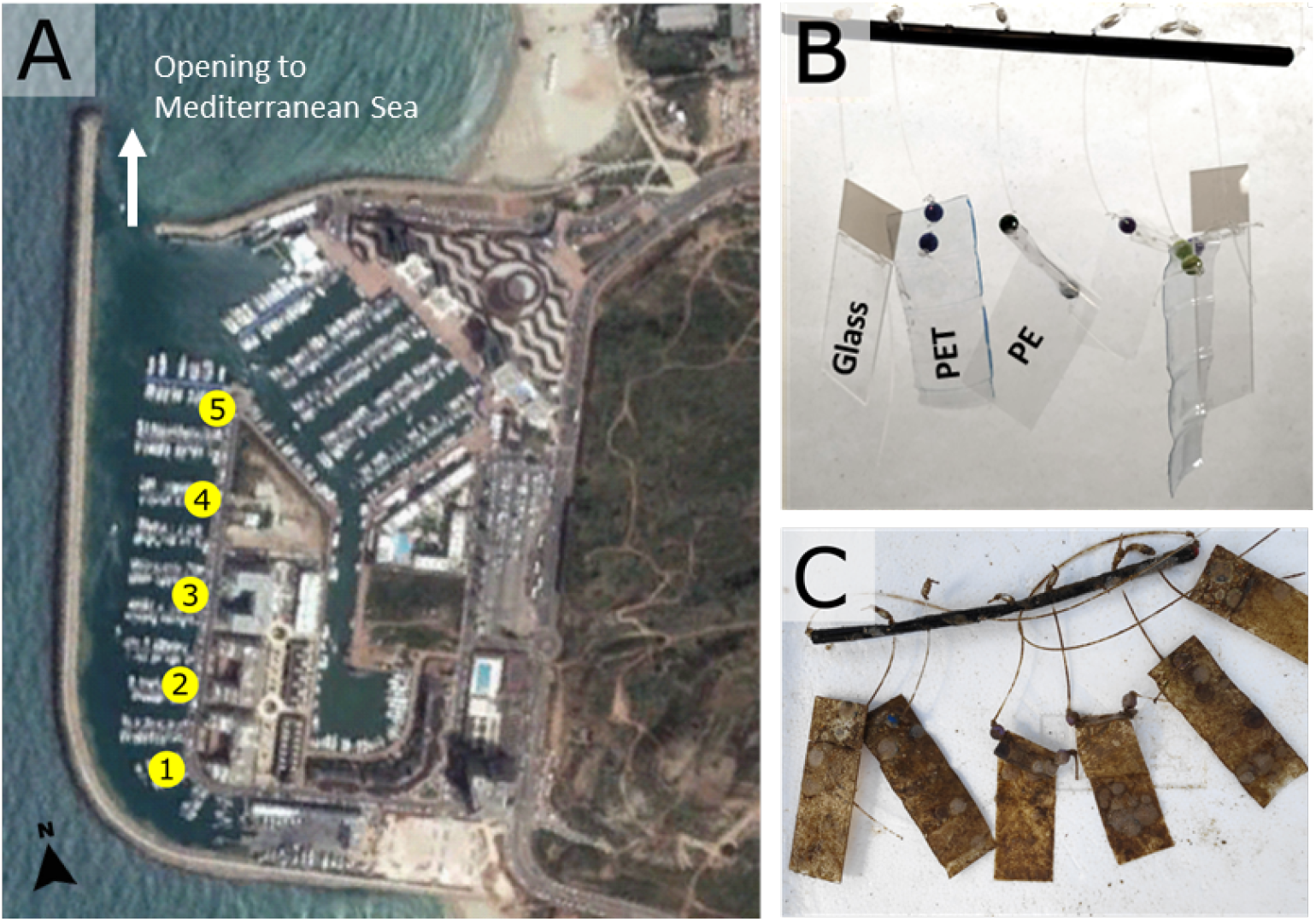
Experiment location and set up. **(A)** Locations of the five replicates that were placed from south to north in Herzliya Marina, Israel (32° 09′ 38.8” N 34° 47′ 35.0” E). **(B)** Each replicate consists of two 17×26 mm flat pieces of each of the three materials (PE, PET and glass) tied with a fishing line to a straw and secured with plastic beads. This ‘mobile’ of materials was secured to the dock such that the surfaces were submerged ∼0.5 m below the surface and 6-8 m above the marina bottom. **(C)** The materials after they were submerged for 30 days in Herzliya Marina water.

### Sample collection and DNA extraction

Each of the mobile materials were sampled in its entirety, gently washed 3 times for 5 min with filtered seawater to remove unbound material and separately processed in each of the subsequent assays and procedures described below. Seawater was sampled in proximity to each of the mobiles (5 times) using sterile sampling bottles. 0.5 L of the sampled water was filtered on 0.22 µm polyethersulfone membrane (Millipore) using a 20 L/min pump (MRC). DNA was extracted using the phenol–chloroform extraction method (Debeljak et al., 2017) (SI-P1).

### PCR amplification and clean up

One sample of each material (water filters, PE, glass and PET) from each mobile (5 repeats) were subjected to PCR amplification with three sets of primers to amplify 3 barcode regions. The complete 16S rRNA gene was amplified using 27F and 1492R primers (Weisburg et al., 1991). Primers 566F and 1289R (Hadziavdic et al., 2014) were used to amplify the V4 and V5 regions of the 18S rRNA gene and ITS86F and ITS4R (Op De Beeck et al., 2014) were used to amplify the fungal ITS2 loci. The amplification parameters and primer details are listed in Table S1. The reaction volume was 50 µL with 25–75 ng of template sample DNA. The PCR products were cleaned with QIAquick-PCR Purification kit (QIAGEN) to meet the criteria of the MinION nanopore library preparation protocol (Karamitros and Magiorkinis, 2018).

### MinION library preparation and multiplexed nanopore sequencing

The sequencing libraries were prepared using the Native barcoding amplicons protocol with EXP-NBD104 and SQK-LSK109 kits (Oxford Nanopore Technologies) according to (Davidov et al., 2020). The 16S and 18S sequencing libraries were loaded to the same MinION flow cell in two batches: the first included the amplified 16S and 18S DNA barcodes of replicates 1, 3 and 5 (12 multiplexed libraries in total), and the second included the amplified 16S and 18S DNA barcodes of replicates 2 and 4 (8 multiplexed libraries in total). The two sequencing runs were separated by a washing step using EXP-WSH002 Kit (Oxford Nanopore Technologies). Each library was loaded onto to the MinION Nanopore Spot-on flow cell (FLO-MIN106D, version R9) and sequenced until reaching ∼7 Giga nucleotides (∼4 M reads). The ITS2 sequencing library was loaded to a new MinION flow cell and sequenced until reaching ∼1.3 Giga nucleotides (∼1 M reads). Base-calling for all libraries were done by the Guppy base calling software 3.3.3, using MinKnow program with the “high accuracy” option. Raw reads were obtained in FAST5 and FASTq formats from which “pass” quality reads were subjected to further analysis.

### Sequence analysis and bioinformatics

Processing and analysis of reads was performed using MetONTIIME pipeline and QIIME2 plugins (Bolyen et al., 2019). Reads were demultiplexed and adaptors and PCR primers were trimmed. Sequences were filtered based on read quality (min_quality; 16S&18S, 9-11; ITS2, 8), and read length was restricted (amplicon_length X, lenfil_tol Y) based on read length histograms, to give the following range: 16S, 1399-1599 nucleotides; 18S, 605-755 nucleotides; ITS2, 100-1100 nucleotides. Clustering and taxonomic classification was performed separately for each barcode. Consensus sequences of 16S rRNA and 18S rRNA genes were mapped against Silva132 database (Quast et al., 2013) with 85% similarity threshold. Consensus sequences of the ITS barcode were mapped against UNITE database V8 (Nilsson et al., 2019) using Blast, with a 70% similarity threshold. The featured tables were imported into R using phyloseq (Team, 2018).

Sequences mapped to chloroplasts, mitochondria, eukaryotes and “unknown organisms” were removed from the 16S analysis, while sequences mapped to bacteria (due to mitochondrial DNA) and “unknown organisms” were removed from the 18S and ITS analysis. Alpha diversity was estimated by richness (observed Operational Taxonomic Units (OTUs)), together with Shannon and Pielou diversity indexes using phyloseq (McMurdie and Holmes, 2013). After alpha diversity analysis, OTUs with a total abundance of 1 were excluded and the remaining OTUs were normalised to relative read abundance (dividing the number of reads for each sample by the total reads count). For Beta diversity we used phyloseq (McMurdie and Holmes, 2013) to perform principle coordinates analysis (PCoA). This ordinated the sequences using the Bray-Curis distance matrix to visualize multivariate structures of the communities. Stacked bar plots and heatmaps were produced using MicrobiomeAnalyst (Chong et al., 2020) and Ampvis2 (Andersen et al., 2018) respectively. We used *limma* (Ritchie et al., 2015) to generate linear models for the detection of differential abundant species between surface materials. Mean location variance was removed from all measurements so the community differences due to surface type could be better seen. We focused on OTUs that had FDR adjusted p. value <0.05 and log2 fold change of 0. Venn diagrams were created with InteractiVenn tool (Heberle et al., 2015), based on OTUs represented by at least two reads. All Nanopore MinION filtered reads analyzed in this project were deposited in the NCBI SRA database(Agarwala et al., 2016) under Bioproject PRJNA680232 (accession numbers: SRX9553811–SRX9553850, SRX10393298-SRX10393305, SRX11261845-SRX11261863).

### Biofilm analysis and microscopy

Scanning electron microscopy (SEM) was used to visualise species and the morphology of the biofilms (SI-P2). For fungal identification, Lactophenol Cotton Blue dye (Sigma-Aldrich) was applied and visualised under a light microscope (SI-P3). A modified crystal violet method (SI-P4) was used for the indirect measurement of the relative biofilm biomass on the experimental samples.

## Results

### High microbial diversity is observed within the one-month-old biofilm

Within 30 days from submersion in the marina waters, all substrates were heavily covered in biofilm (Fig. 1C). The biomass was similar on all surfaces, as indicated by the crystal violet assay (Fig. S1). SEM imaging revealed that the plastic surfaces were mostly covered with bacterial biofilm (Fig. 2A), exposed bacteria (Fig. 2B) and multiple diatom species (Fig. 2C-E). We were also able to identify dinoflagellates (Fig. 2E), bryozoans (Fig. 2F) and fungal-like fruiting bodies (Fig. 2G). Furthermore, we observed multicellular filaments (Fig. 2H), of which some were identified as algae based on the presence of chlorophyl and some were confirmed to be fungi hyphae by Lactophenol Cotton Blue staining (Fig. 2I).

**Figure 2.**
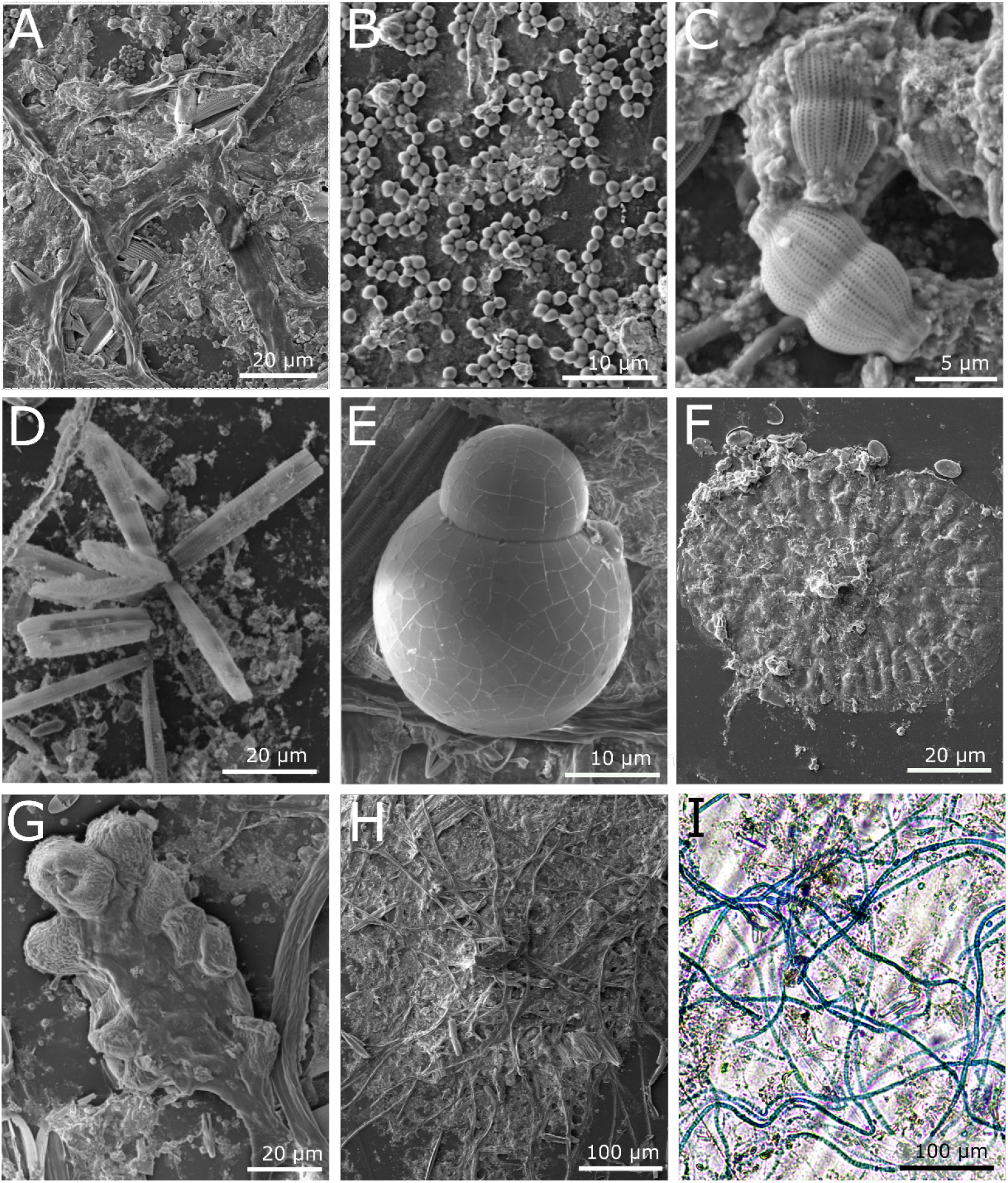
Microscopy images of the microbial community on plastic surfaces from marine waters after 1 month. **(A-H)** SEM images; **(A)** Biological coverage on plastic surface after 1 month, **(B)** bacterial colonies on the plastic surface, **(C-D)** diatoms associated with biofilm, **(C)** *Fragilariopsis sp*. **(D)** *Amphora sp*., **(E)** a dinoflagellate, **(F)** Bryozoan, **(G)** fruiting body, **(H)** multicellular filaments. **(I)** Light microscopy image of a network of fungal hyphae specifically stained with Lactophenol Cotton Blue.

The nanopore sequencing run produced an average of 25235 16S reads, 80125 18S reads, and 24777 ITS reads per sample. The average read length was 1428 nucleotides for 16S, 669 for 18S and 369 for the ITS barcode. However, while both 16S and 18S ribosomal amplicons had read length standard deviation of 24 and 20 accordingly, the ITS reads varied greatly in length with standard deviation of 154.5 nucleotides. The mapping rates of the ITS sequences were very low (3% on average) compared to the 16S and 18S mapping rates (both 99%), and the ITS sequences had a lower taxonomic resolution (Table S2).

Community complexity parameters including richness (number of observed OTUs), evenness (Pielou’s index) and diversity (Shannon’s index) were obtained for each of the sample types (water, PET, PE and glass) based on 16S and 18S reads (Fig. S2). For 16S sequence analyses, the water had significantly lower taxonomic evenness, contributing to lower OTU diversity (Shannons index) in comparison to the surface samples (Fig. S2A). This agrees with previous environmental studies that showed the same trend (De Tender et al., 2015; Bryant et al., 2016; Didier et al., 2017). In contrary, the 18S analysis resulted in no significant difference among the samples in any of the diversity indexes (Fig. S2B). Previous 18S-based analyses report water communities as more diverse (Didier et al., 2017; Kettner et al., 2019; Dudek et al., 2020). Because of the low ITS mapping rates, we did not analyse the alpha diversity parameters for fungi.

To assess the similarities in the community composition between the samples, beta diversity analysis was performed using PCoA analysis (Fig. 3). In both 16S and 18S analyses, the five water samples formed distinctive clusters from the surface samples (Fig. 3A-D). When excluding the water samples, no distinct clusters were observed in any of the analyses (Fig. 3B-F). In contrast, in the ITS analysis, while the water samples showed homogeneity among themselves, they did not form a separate cluster from the other samples (Fig. 3E-F). This result suggests overlapping surface-bound and free-living fungal taxa distribution.

**Figure 3.**
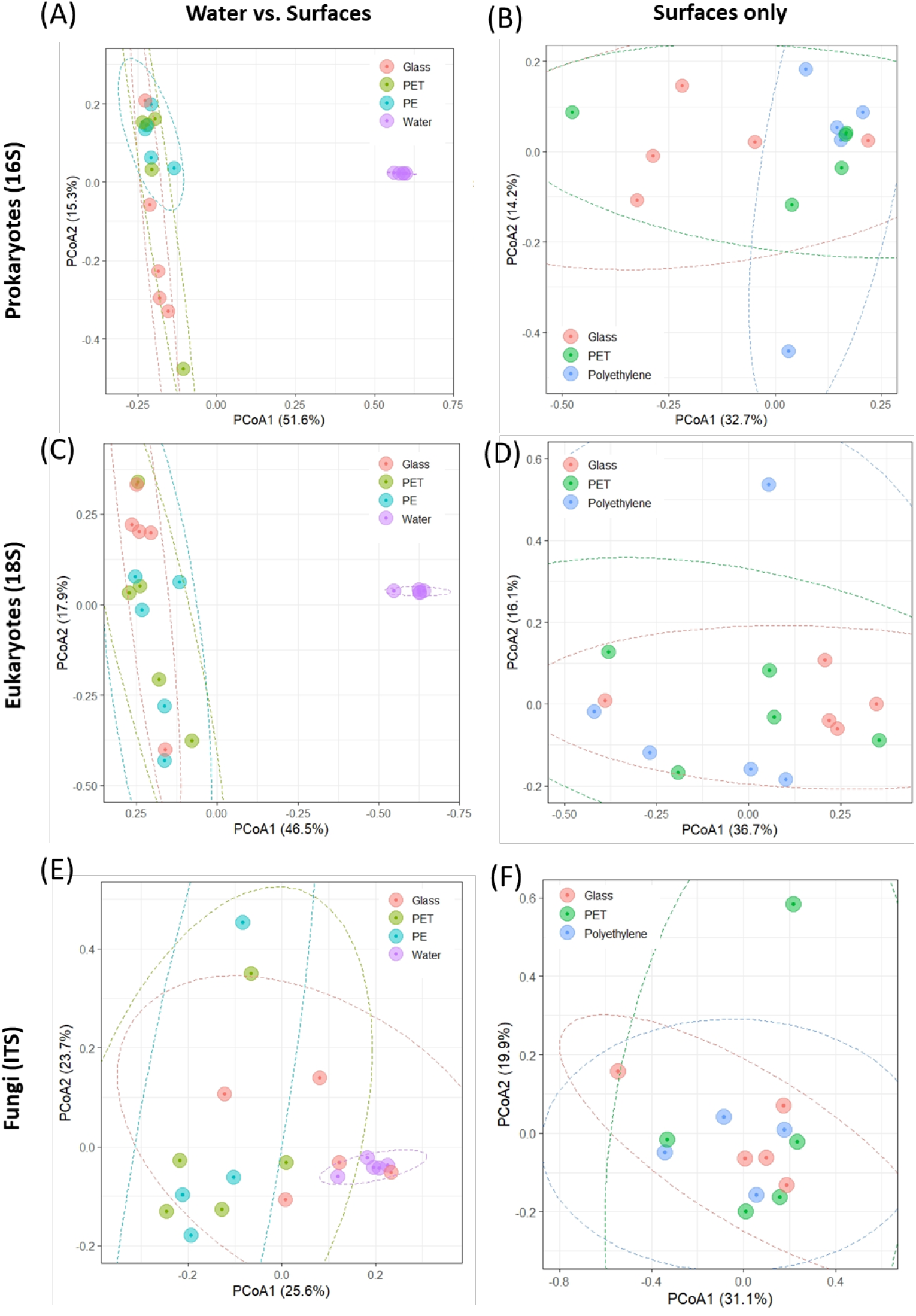
PCoA of the Bray-Curtis dissimilarities among the plastisphere communities of the samples (N=5). **(A-B)** 16S microbiomes with the water community (left) or without it (right). **(C-D)** 18S microbiomes with the water community (left) or without it (right). **(E-F)** ITS microbiomes with the water community (left) or without it (right).

### Substrate specificity of the 16S microbial communities

The filtered, high quality 16S sequences were clustered into an average of 2733 operational taxonomic units (OTUs) for PET, 21129 for PE and 4163 for glass, corresponding to 524, 754 and 677 taxa respectively. The top 10 most abundant prokaryotic genera, that were identified based on the mapped read counts, contain some different genera across the surfaces (Fig. 4A). *Candidatus, Ekhidna, Muricauda* and *Portibacter* were included in the top 10 most abundant genera on PE and not in the other surfaces, whereas *Rhodopirellula* and OM60 clade were included only in the PET top 10. Among the above genera, *Muricauda, Candidatus* and *Rhodopirellula* have been previously reported as hydrocarbon-degrading bacteria (Jiménez et al., 2011; Didier et al., 2017; Laso-Pérez et al., 2019; de Araujo et al., 2021).

**Figure 4.**
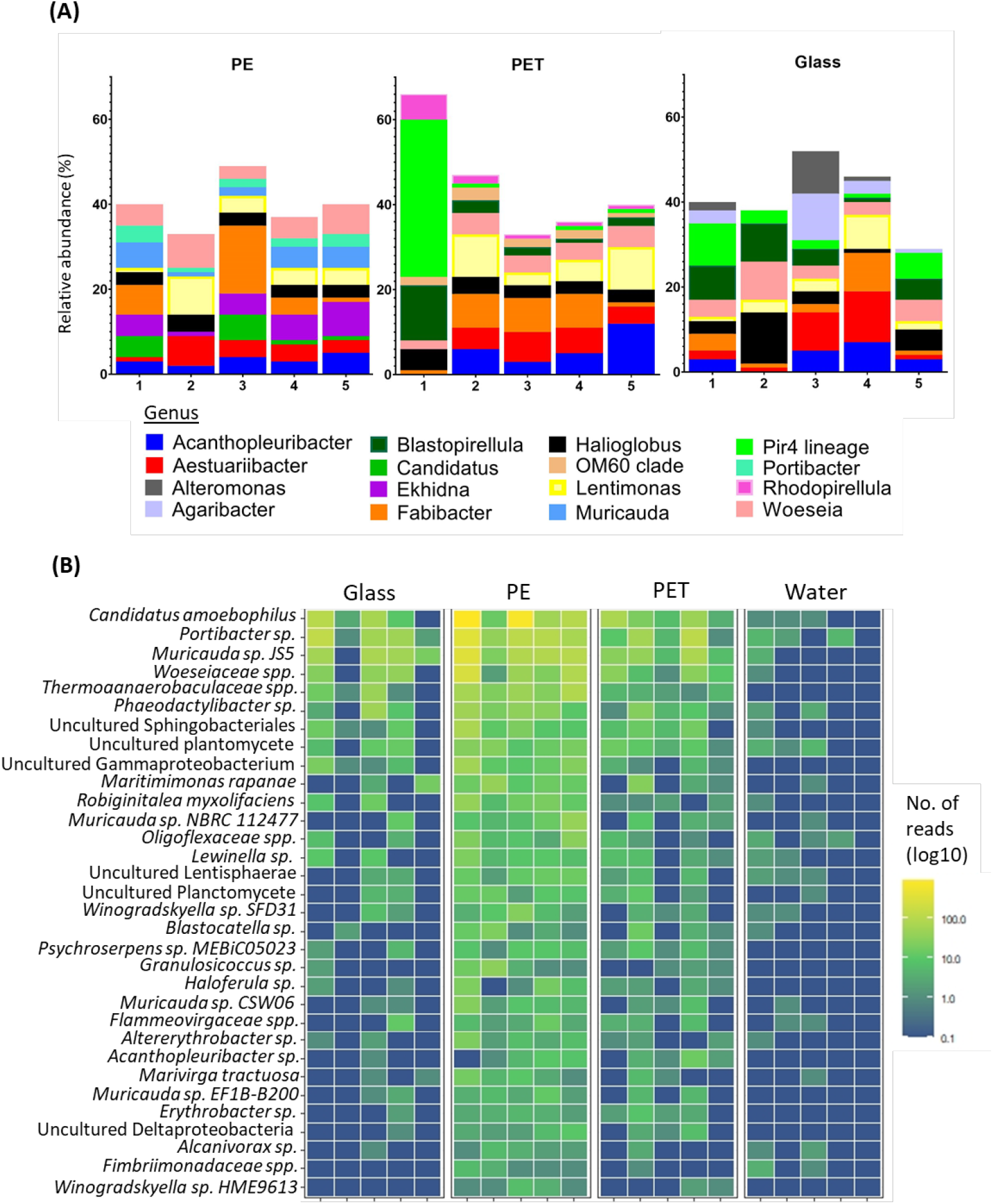
Substrate specificity of bacteria based on 16S mapped reads. **(A)** Relative abundance of top 10 abundant genera. **(B)** Heatmap of the bacterial taxa with higher mapped read ratios in the PE samples compared to glass and water samples, across all replicates.

To identify species with preference to plastic, we searched for species with significantly higher read ratios in the PET and PE samples as opposed to glass. Linear discriminant analysis of our 16S rRNA gene sequences revealed 35 OTUs, corresponding to 32 taxa that were significantly more abundant in the PE samples than in the glass samples (Fig. 4B) (Fig. S3). Of the 32 taxa, 12 were resolved at the species level and 10 at the genera level (leaving 11 of higher taxonomic classification). Comparisons between PET and Glass or PET and PE did not result in the identification of differentially abundant species.

Among the bacterial species that showed significantly higher read representation in the PE samples compared with the glass and water samples, *Candidatus Amoebophilus* showed highest significance values (adjusted p. value = 0.003) (Table S3). We also identified 4 species of the genus *Muricauda*, which matches with the top 10 genera for PE (Fig. 4A), and 2 species from the genus *Winogradskyella*, which has been previously identified as a hydrocarbon degrading genera (Wang et al., 2014). Other species that were significantly differentially abundant on PE included *Maritimimonas rapanae, Robiginitalea myxolifaciens* and *Psychroserpens sp. MEBiC05023*, all from the *Flavobacteriaceae* family, and *Marivirga tractuosa*. Of the unresolved taxa, *Alcanivorax sp*. is a well-known degrader of alkanes and petroleum (Yakimov et al., 2019). It was also recently shown that certain *Alcanivorax* species are able to degrade PE (Delacuvellerie et al., 2019).

### Substrate specificity of the 18S eukaryotic communities

The high quality 18S sequences were clustered into an average of 10997 OTUs for PET, 11608 for PE and 7956 for glass, corresponding to 730, 735 and 506 taxa respectively (Table S2). Within the 18S mapped reads, the genera with the highest mapped read proportions across all surfaces was the bryozoan *Amathia*. However, there were also differences between the top 10 most abundant genera among the surfaces (Fig. 5A). On PE, the most abundant genera included *Scytosiphon*, a genus of brown seaweed, the bryozoan *Bugulina*, the entoprocta *Barentsia* and the sabellid polychaete *Parasabella*. On the other hand, the top 10 genera on PET included the copepods genus *Acartia*, as well as the diatom genus *Nitzschia* and the ciliate genus *Dysteria*.

**Figure 5.**
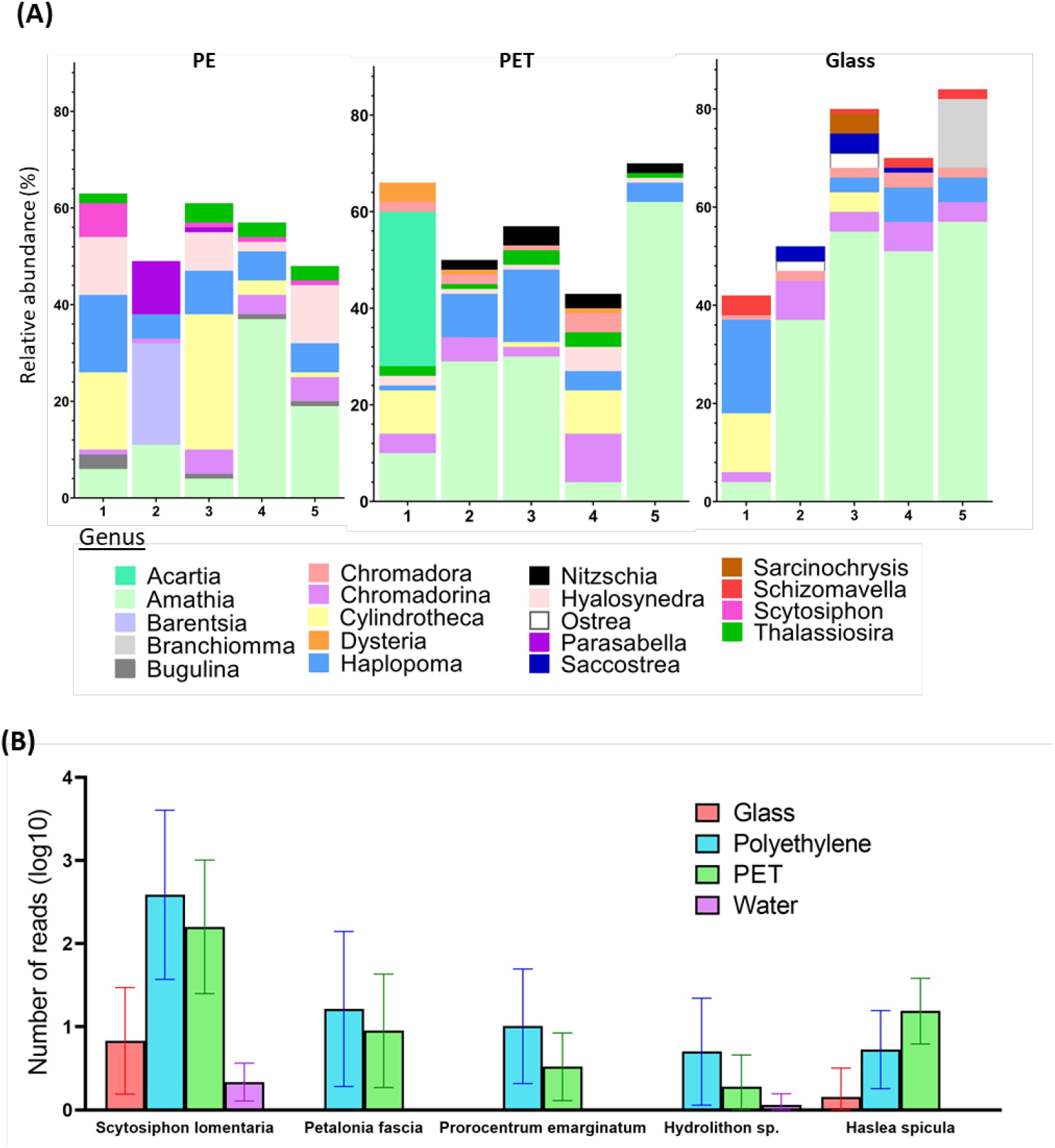
Substrate specificity of eukaryotes at the genus and species levels based on 18S. **(A)** Relative abundance of top 10 abundant genera based on metabarcoding with 18S. **(B)** Eukaryotic taxa with differential mapped read counts in the plastic samples compared to the glass samples, across all replicates.

Linear discriminant analysis of our 18S rRNA gene sequences identified five OTUs, corresponding to four species and one genus, with significantly higher read ratios in the plastic samples compared to glass samples (Table S4). These species included two brown algae; *Petalonia fascia* and *Scytosiphon lomentaria*, with higher read ratios in both PET and PE samples. The benthic dinoflagellate *Prorocentrum emarginatum* and the red alga *Hydrolithon sp*. had higher read representation in the PE samples, while reads mapped to the diatom *Haslea specula* were found in higher ratios in the PET samples (Fig. 5B).

### Marine fungi communities on plastic surfaces

For the identification of fungal taxa we used the ITS2 barcode. The ITS2 locus has been shown to be the most suitable taxonomic barcode for the characterization of fungal communities and has been previously used for the identification of fungi from marine plastic debris (Op De Beeck et al., 2014; Davidov et al., 2020).

Due to the limitations of the available databases for marine fungi, only 3% of the reads were mapped to reference sequences. Despite this, our analysis still identified an average of 57 fungal OTUs in the PET samples and 58 in the PE samples, of which 23 and 25 respectively were classified to the species level. We also identified an average of 87 fungal OTUs in the glass samples and 114 in the water samples, of which 29 and 41 respectively were classified to the species level (Table S2).

In contrast to the 16S-and 18S-based beta-diversity analyses, the fungi (barcoded with ITS) analysis did not show distinction between the taxonomic composition of the water vs. the composition of the surface biotas (Fig. 3E). However, within the ITS mapped reads, there were still differences between the top 10 abundant genera across the surfaces (Fig. 5A). On PE, the most abundant genera included *Neocatenulostroma, Penicillium* and *Vishniacozyma*. Whereas in the top 10 genera of the PET samples we identified *Candida, Cyberlindnera* and *Rhodosporidiobolus*. Due to the limited mapping results, differential abundance analysis was not performed. Nevertheless, of all 316 OTUs identified, five were found only on PE (Fig. 6B) corresponding to species *Aspergillus penicillioides, Bipolaris sorokiniana, Filobasidium magnum, Knufia mediterranea* and *Ramichloridium cucurbitae*. Another five OTUs were uniquely found on the PET samples corresponding to *Cryptococcus aspenensis, Cyberlindnera jadinii, Debaryomyces vindobonensis, Pyrenochaetopsis leptospora*, and *Symmetrospora coprosmae*. Another two OTUs corresponding to the species *Candida sake*, and *Peniophora lycii* were identified in the PET and the PE samples but not in the glass samples and the water.

**Figure 6.**
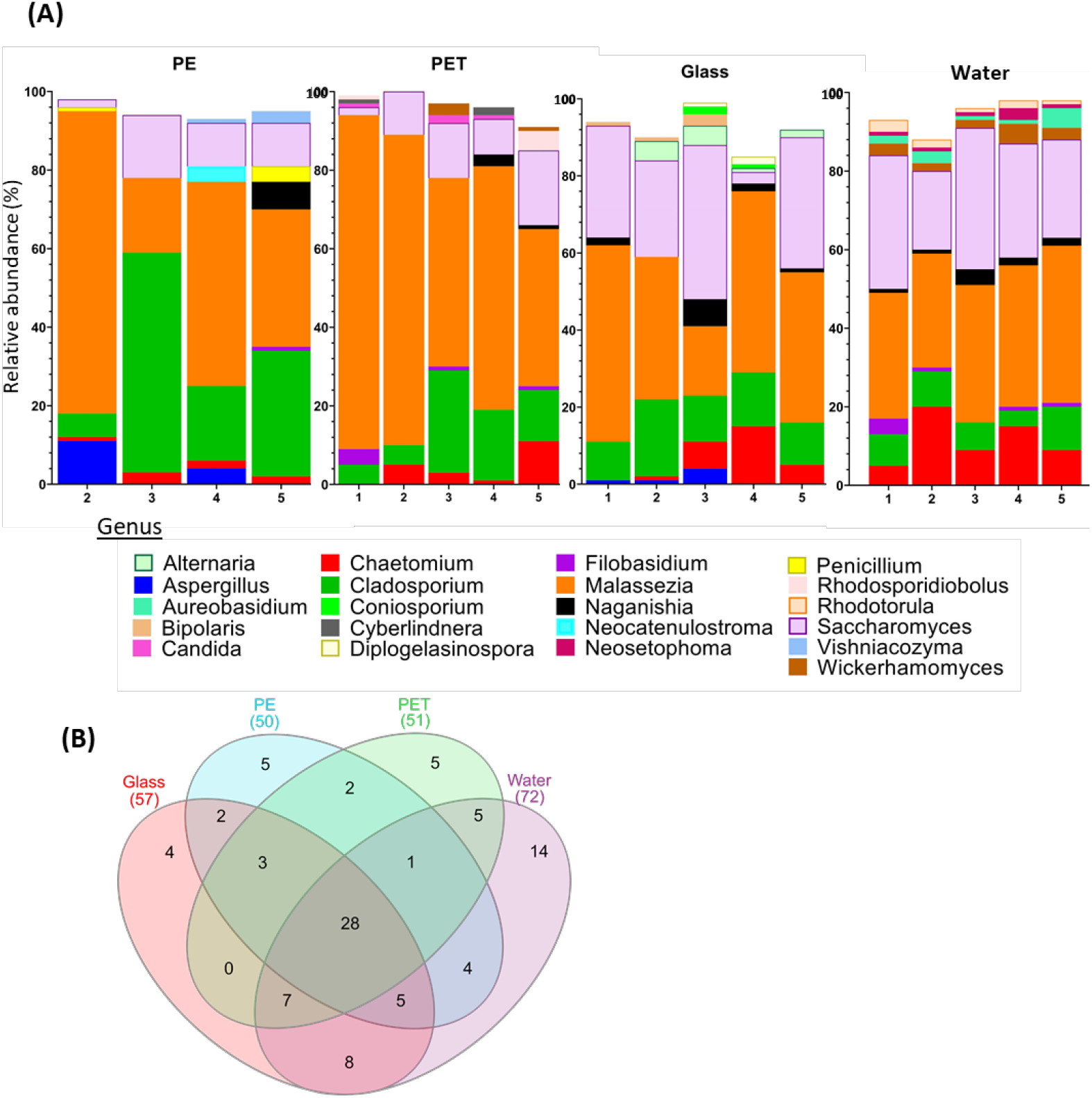
ITS fungal metabarcoding analysis. **(A)** Top 10 abundant fungi genera based on ITS metabarcoding. **(B)** Shared and unique fungal OTUs among the surfaces and the water.

## Discussion

In this study we analysed one-month-old Mediterranean Sea plastisphere communities and screened for prokaryotes and eukaryotes species, including fungi, with differential abundance on PET and PE plastic surfaces compared to glass. Many studies have focused on the search for plastic-specific taxa (reviewed in (Latva et al., 2021)). However, very few resulted in the identification of the taxonomic levels of genera and species. For achieving a higher taxonomic resolution, including the species level, we used longer barcode regions that were sequenced with the nanopore MinION platform. Additionally, we characterized the composition of the fungal biome using a shorter, well-established barcode within the ITS2 locus.

Previous 16S and 18S metabarcoding analyses have repeatedly demonstrated that the plastisphere is a separate ecological niche from the surrounding water (Zettler et al., 2013; Dussud et al., 2018; Frère et al., 2018). Our metabarcoding beta diversity analysis supports this observation. On the other hand, this pattern was not repeated in the ITS metabarcoding analysis of fungi. Instead, it showed a taxonomic overlap between the surface-bound and free-living fungal taxa. This overlap was previously explained by the ability of some fungal species to display amphibious qualities (Velez et al., 2015; Amend et al., 2019) and based on possible symbiotic relationships with other marine organisms (Redman Regina et al., 2001; Amend et al., 2019; Gladfelter et al., 2019) that may be present in both environments. Another possible explanation is the presence of reproduction products of the surface-bound fungi in the water (Lacerda et al., 2020). We would like to note that a definitive conclusion cannot be drawn for this kingdom because of the low mapping rates of the ITS sequences (∼3%). Other studies that have used the ITS barcode resulted in similarly low mapping rates of 0.01 to 4% (Kettner et al., 2017; Lacerda et al., 2020; Wang et al., 2021) which is attributed to the limited taxonomic coverage of the available fungal databases, especially when it comes to marine fungi (Hawksworth and Lücking, 2017).

So far, most environmental studies did not find conclusive differences between the microbial communities on plastic vs. glass surfaces (Oberbeckmann et al., 2016; Pinto et al., 2019; Erni-Cassola et al., 2020; Laverty et al., 2020; Zhao et al., 2021). On the other hand, studies in enclosed or semi-enclosed systems often showed differential taxonomic representation (Kirstein et al., 2018, 2019; Ogonowski et al., 2018; Pinto et al., 2020). These differences between the two types of studies may be due to the milder masking effects of environmental factors under controlled lab conditions. For this reason, we find that the location of our experiment, within the semi-protected environment of the marina that is connected to the open sea, may serve as an ideal location for such comparative studies of the natural environment.

To identify the high-resolution differences in the taxonomic composition of the microbiomes, we first analysed the top 10 most abundant genera in each surface. This analysis resulted in several genera that were recurrently identified on one substrate and not in the other. However, it has been established that the OTUs contributing to the most dissimilarity between substrates are not necessarily the most abundant ones (e.g. (Kirstein et al., 2018, 2019)). Therefore, it is important to use the appropriate statistical tools to identify differentially abundant species. Here we took advantage of the *limma* package, which implements statistical algorithms developed for the analysis of differential expressed genes. These specialised algorithms make statistical conclusions more reliable when the number of samples are small and have different levels of variability and complex set ups (Ritchie et al., 2015). Our analysis identified 32 prokaryotic taxa for which the mapped 16S reads were significantly differentially abundant in the PE samples compared to the glass samples. Many of which belong to genera that have been previously reported in association with plastic communities including, *Maritimimonas* (De Tender et al., 2017), *Saprospiraceae, Flammeovirgaceae* and *Lewinella* (reviewed in (Roager and Sonnenschein, 2019)) as well as *Fulvivirga* (Tourova et al., 2020) and *Cyclobacteriaceae* (Miao et al., 2019). Moreover, our differential abundance analysis identified bacteria taxa that are considered to be hydrocarbon and plastic-degrading including genera *Muricauda* (Didier et al., 2017), *Winogradskyella* (Wang et al., 2014) and *Alcanivorax* (Delacuvellerie et al., 2019). The same analysis for the 18S sequences identified five taxa that were differentially abundant on PE and PET compared with glass including two species of brown algae, *Scytosiphon lomentaria* and *Petalonia fascia* which was previously identified in the plastic microbiome (Ibabe et al., 2020). The dinoflagellate *P. emarginatum* and the red algae *Hydrolithon sp*., that had significantly different read ratios on PE, have also been reported to dominate plastisphere communities (Dudek et al., 2020). The diatom *Haslea spicula*, which was significantly differentially abundant on the PET surfaces is a mobile pennate diatom that is known to colonise artificial surfaces (Winfield et al., 2018). The presence of diatoms on marine plastic has been repeatedly shown (e.g. (Oberbeckmann et al., 2014; Reisser et al., 2014; Davidov et al., 2020; Dudek et al., 2020)) and was clearly observed in our SEM and light microscopy imaging.

The fungal biome on marine plastic has been so far understudied and only a handful have analysed the ITS genes (e.g. (Zhang et al., 2015; De Tender et al., 2017; Davidov et al., 2020; Lacerda et al., 2020)). Our ITS metabarcoding analyses and lactophenol cotton blue staining showed the plastic surfaces in the marina were colonized by a highly developed network of fungi, mostly of the genera *Malassezia (*phylum Basidiomycota), *Cladosporium* and *Saccharomyces* (phylum Ascomycota). The presence of *Cladosporium* on plastic debris was previously shown (De Tender et al., 2017; Lacerda et al., 2020; Xue et al., 2021). 12 fungal OTUs were found to be unique to the plastic samples including the species *Pyrenochaetopsis leptospora, Candida sake, Debaryomyces vindobonensis, Aspergillus penicillioides* and *Bipolaris sorokiniana*, a wheat pathogen that causes leaf spot disease (Ye et al., 2019). *Peniophora lycii*, a species that was found on both PE and PET, but not on glass or water, has recently been shown to secrete three laccase isozymes (Glazunova et al., 2020), that may be capable of breaking down non-hydrolysable plastics such as PE (Inderthal et al., 2021). Many marine fungi can degrade complex hydrocarbons (reviewed in (Amend et al., 2019)) and often dominate in oil polluted environments (Bik et al., 2012; McGenity et al., 2012). So far only one marine fungi, *Zalerion maritimum* has been found to degrade plastic (PE) in laboratory conditions (Paço et al., 2017; Santacruz-Juárez et al., 2021). Fungi are an abundant and active component of the ocean environment with plastic degradation potential that warrants further investigation.

While DNA metabarcoding of the plastisphere biome continues to be a favorable approach for its taxonomic composition characterization it needs to be fine-tuned to be effective in the identification of plastic-specific genera and species. Refining the resolution and the scope of this approach will provide useful information that can be the basis for species-targeted studies to unveil the molecular mechanism for plastic colonization and metabolism.

## Acknowledgements

The authors are grateful to Natali Litvak for the SEM imaging, to Iryna Yakovenko for her help in the field and lab work and to Dr. Maxim Rubin-Blum for scientific advice. This work was supported by the Israel Ministry of Science and Technology Israel-Portugal collaboration Grant 3-16503.

## Notes

### Competing Interest Statement

The authors have declared no competing interest.

