## Supplementary information for "High-resolution screening for marine prokaryotic and eukaryotic taxa with selective preference for PE and PET surfaces"

**Supporting Information** (11 pages)

SI-P1. DNA extraction………………………………………………………………………………………………………S
SI-P2. Scanning Electron Microscopy………………………………………………………………………………S
SI-P3. Light microscopy, lactophenol blue staining…………………………………………………………S
SI-P4. Biofilm quantification by crystal violet method…………………………………………………….S
Figure S1. Biofilm quantification by crystal violet method………………………………………………S
Table S4. Differentially abundant 18S OTUs……………………………………………………………………S
Figure S4. Differential abundant 18S OTUs between different samples…………………………S
### Table S1. Primer and amplification details*

| **Gene/genetic locus** | **Primers** | **Primer sequences (5' -> 3')** | **Annealing temp.** | **Product Length (bp)** |
| --- | --- | --- | --- | --- |
| 16S-small subunit ribosomal RNA | 27-F 1492-R | AGAGTTTGATCMTGGCTCAG GGTTACCTTGTTACGACTT | 56°C | 1499 |
| 18S-small subunit ribosomal RNA | 566-F 1289-R | CAGCAGCCGCGGTAATTCC ACTAAGAACGGCCATGCACC | 57°C | 723 |
| ITS2- second nuclear ribosomal internal transcribed spacer | ITS86-F ITS4-R | GTGAATCATCGAATCTTTGAA TCCTCCGCTTATTGATATGC | 55°C | 369 |

* PCR parameters: 2 minutes at 94°C ,32 cycles of: 30 seconds at 94°C, 30-90 seconds at 45°C to 57°C, 136 30-90 seconds at 72°C, and final extension 72°C for 5 minutes.

**SI-P1** DNA extraction

DNA was extracted using the phenol–chloroform extraction method. The samples (water filters, PE, glass and PET pieces) were immediately put into a sterile sampling bag with 10 mL of lysis buffer (10 mM Tris–HCL pH8, 25 mM Na_2_ EDTA pH8, 1v/v% SDS and 100 mM NaCl), subjected to manual squeezing and processed and stored in −20 °C until processing. Samples were later thawed, subjected to bead beating with ~ 0.4 g of 425–600 µm sterile glass beads (Sigma) and proteinase K (5 units/µL) and Lysozyme (2000 units/µL) digestion. All other steps of the DNA extraction were performed according to ^25^. DNA was finally eluted in 40 µL EB (10 mM TE Tris 1 mM EDTA pH8).

**SI-P2** SEM- Sample preparation, imaging

Microorganisms on the plastic surfaces were visualised with scanning electron microscopy (SEM) according to ^19^. PE samples were fixed for 2–5 h in 1% glutaraldehyde and 4% PFA for one hour at room temperature followed by washes (three times for 5 min) in distilled water. Samples were kept in 50% ethanol in phosphate-buffered saline (PBS) at − 20 °C until use. One day before use, samples were dehydrated in graded ethanol series for 10 min each in 50%, 70%, 85%, 95% ethanol, followed by 3 ׳ 15 min in 100% ethanol. Dehydrated plastic samples were air-dried for at least five hours in a hood, sputter-coated with 10 nm of platinum/gold (Quorum Q150T ES) and then visualized and imaged on a high-resolution SEM (HRSEM) (Tescan, MAIA 3, TESCAN Ltd., Brno, Czech Republic) in a voltage range of 3-7 kV.

**SI-P3** Light microscopy - Lactophenol Cotton Blue staining, imaging

Light microscopy (LM) was performed using Nikon eclipse Ci-L light microscope fitted with Nikon DS-Fi3 High-definition camera (CMOS). Color corrections, scale and size measurements were performed using ImageJ. For fungal identification, 1-2 drops of Lactophenol Cotton Blue dye (Sigma-Aldrich) were applied on the dry PE samples after fixation with 4% PFA. Stain was washed using 1xPBS until all residual dye was removed. For imaging, samples were positioned on microscope glass slide with a coverslip.

**SI-P4** Biofilm quantification by Crystal violet method

Biofilms were quantified according to the method described by ^40^. At room temperature, each surface material (1 ×2 cm, n=5) was washed with artificial seawater and air-dried in sterile Petri dishes for 45 min. 15 ml of crystal violet (0.1% w/v in artificial seawater) was added to each surface material and incubated for 45 minutes on a shaker. The stained samples were washed three times with artificial seawater until the attached biofilm was clearly visible. Stained samples were air-dried for another 45 min and transferred to a new petri dish containing 15 mL ethanol (95% v/v) and inverted to mix. After 10 min, absorbance of the solution was measured at 595 nm with a nanophotometer (IMPLEN).

**Figure S1**. Biofilm quantification by crystal violet method.

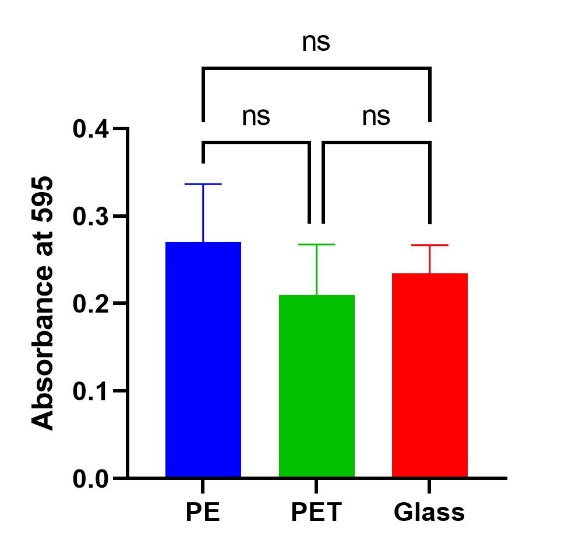

Table S2**.** Minion sequencing results of microbial communities on water, PET, PE, glass using 16S, 18S and ITS primers.

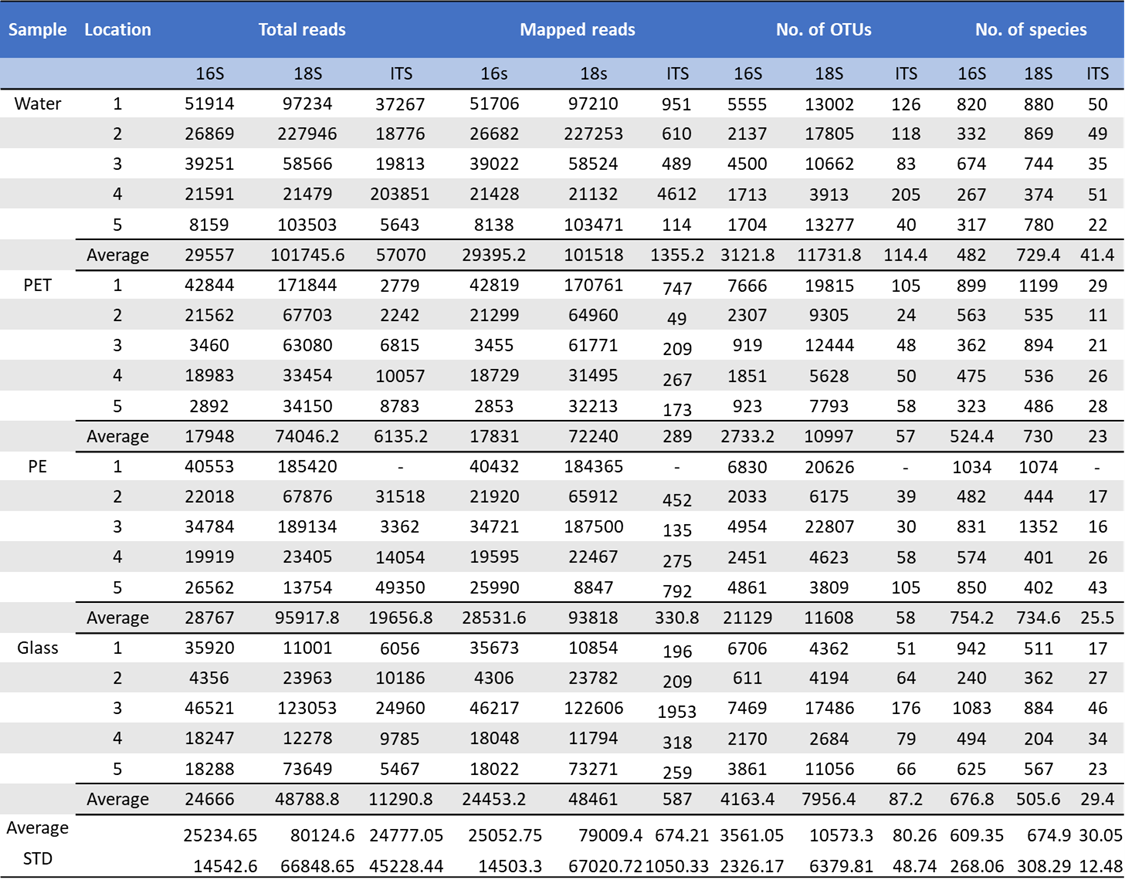

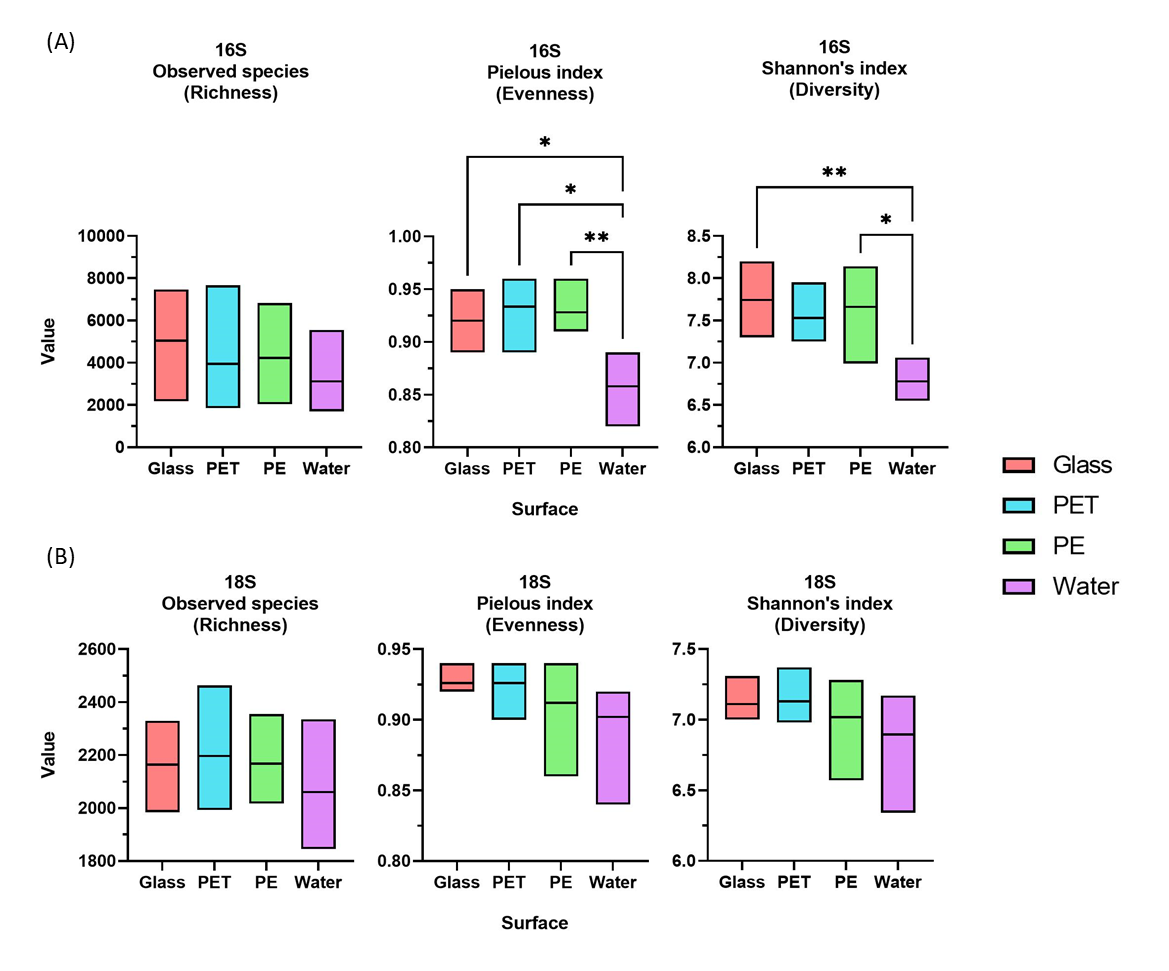
Figure S2. Richness, evenness and diversity of (A) prokaryotic (16S) communities, (B) eukaryotic communities (18S), N =5. Box extends from the minimum to the maximum and line represents mean. Significance was assessed using one-way ANOVA and Tukeys multiple comparison test, *= p<0.05, **= p<0.01. Three samples with less than 5,000 reads (16S PET #3, PET #5 and Glass #2;) were excluded from the alpha diversity analysis.

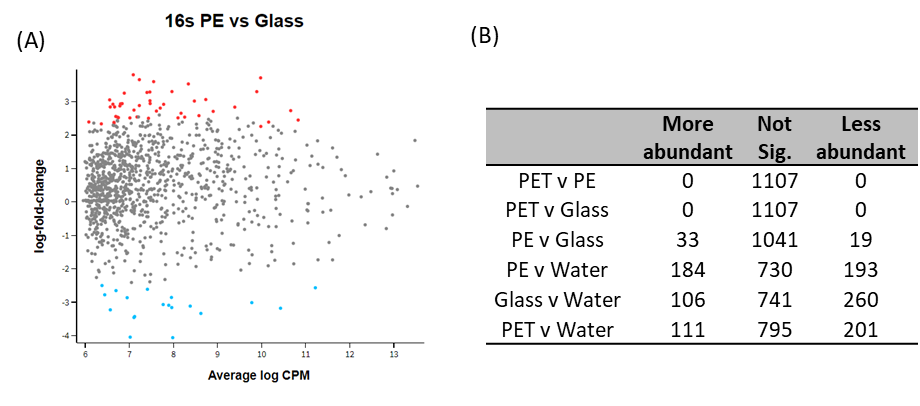

Figure S3. 16S OTUs with a preference to plastic over glass. (A-B) The data was produced based on the ratios of mapped 16S reads. (A) Volcano plot shows the spread of differential abundant 16S OTUs. Each dot represents an OTU, red dots represent OTUs significantly more abundant on PE and blue dots are significantly more abundant on glass. Adjusted p-value = 0.05. (B) Numbers of differential abundant OTUs between different samples.
Table S3. Complete list of significantly differently (positive and negative) abundant 16S rRNA taxa between PE and Glass. Ordered with most significant first (lowest adjusted p-value). Also shown is log counts per million, log fold change and adjusted p-value. Cut off for significance is an adjusted p-value of 0.05.

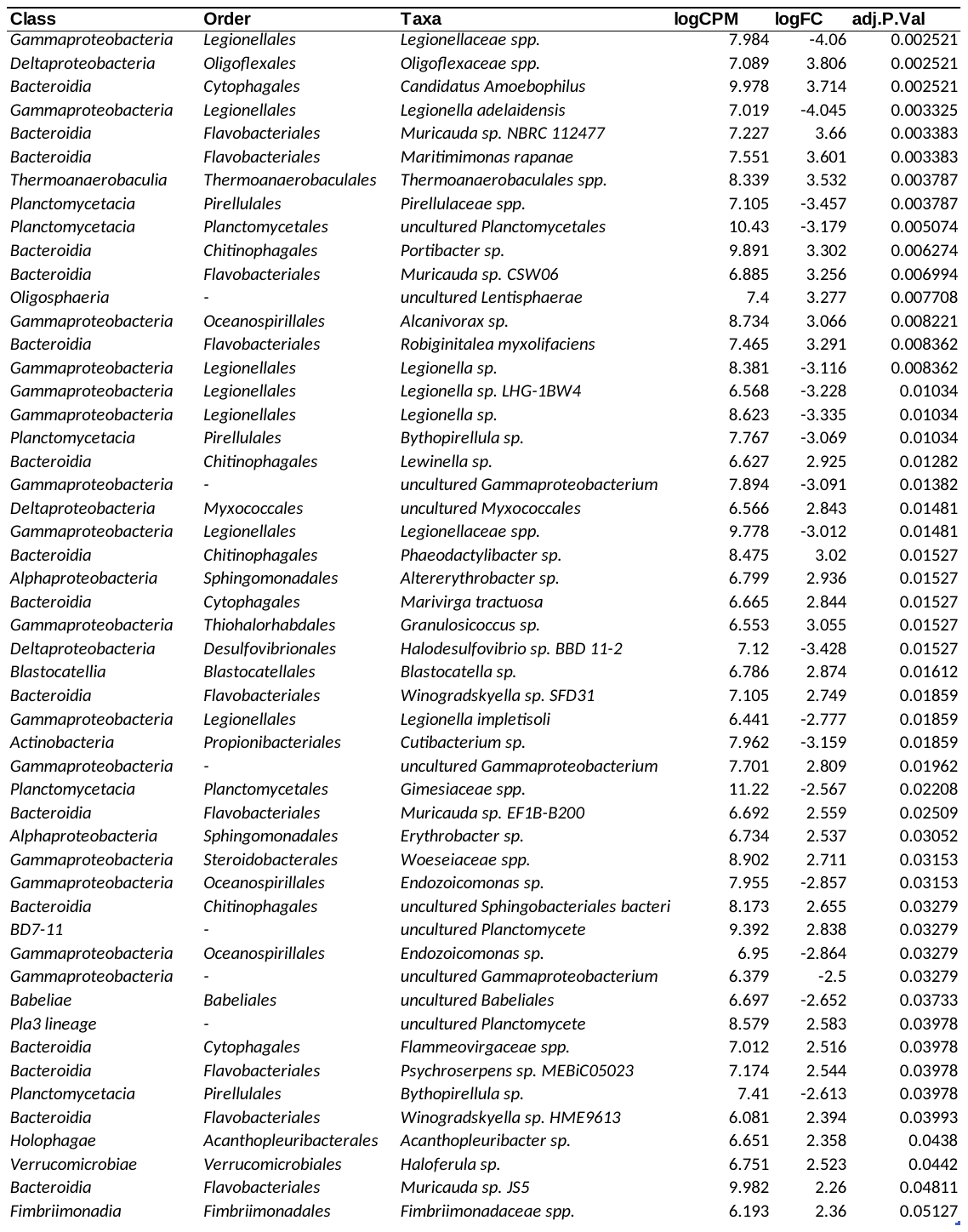

Table S4. 18S rRNA taxa that are significantly more abundant on (A) PE in comparison to glass and (B) PET in comparison to glass with log counts per million, log fold change and adjusted P value.

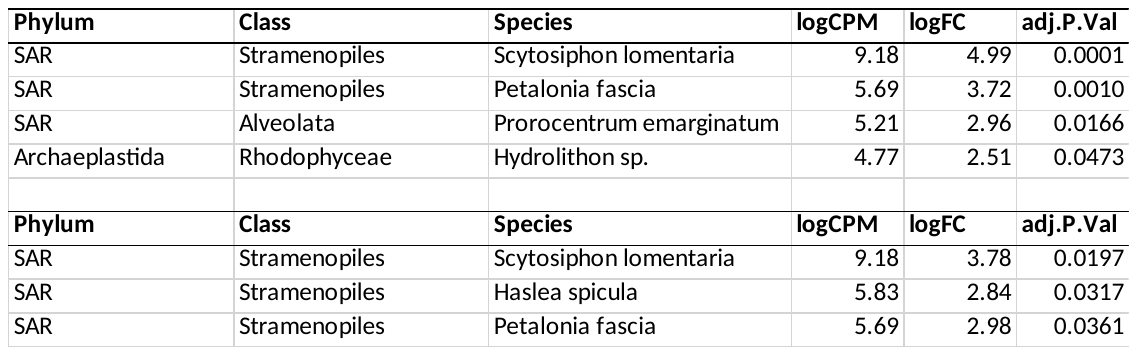

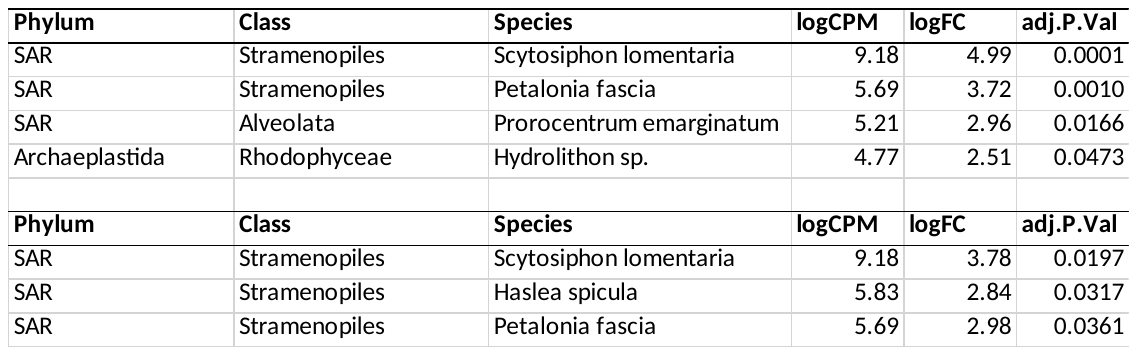

(A)

(B)

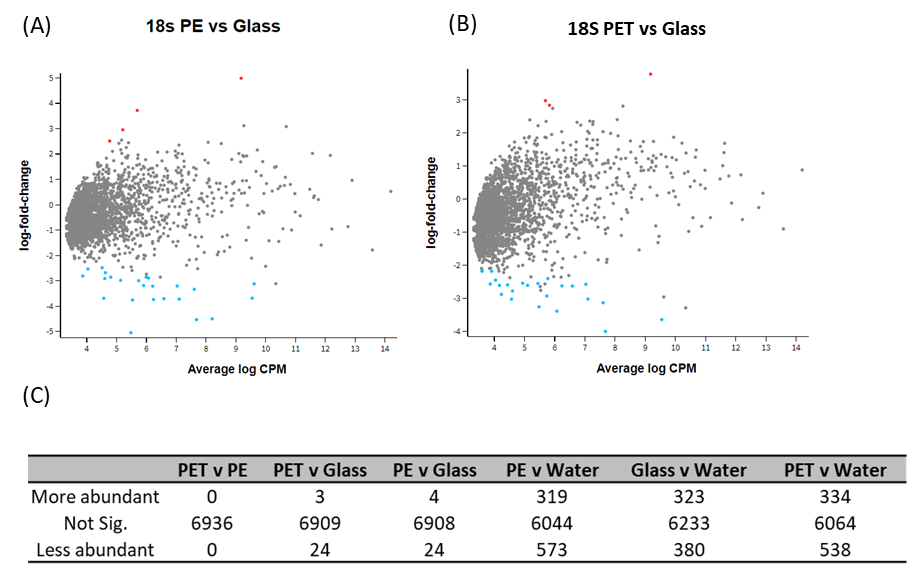
**Figure S4.** 18S OTUs with a preference to plastic over glass. The data was produced based on the ratios of mapped 18S reads. Volcano plot shows the spread of differential abundant OTUs between (A) PE and Glass and (B) PET and Glass. Each dot represents an OTU, red dots represent OTUs significantly more abundant on PE/PET and blue dots represent OTUs significantly more abundant on glass. Adjusted p-value = 0.05. (C) Numbers of differentially abundant OTUs between different samples.
